# Brain structural correlates of autistic traits across the diagnostic divide: A grey matter and white matter microstructure study

**DOI:** 10.1101/2021.05.29.446259

**Authors:** Varun Arunachalam Chandran, Christos Pliatsikas, Janina Neufeld, Garret O’Connell, Anthony Haffey, Vincent DeLuca, Bhismadev Chakrabarti

**Affiliations:** Centre for Autism, School of Psychology and Clinical Language Sciences (SPCLS), University of Reading, UK; School of Psychology and Clinical Language Sciences, University of Reading, Harry Pitt Building, Earley Gate, Whiteknights Road, Reading, RG6 6AL, UK; Centro de Ciencia Cognitiva, Facultad de Lenguas y Educación, Universidad Antonio de Nebrija, Calle de Sta. Cruz de Marcenado, 27, 28015, Madrid, Spain; Center of Neurodevelopmental Disorders (KIND), Centre for Psychiatry Research, Department of Women’s and Children’s Health, Karolinska Institutet & Stockholm Health Care Services, Stockholm.; School of Mind and Brain, Humboldt University, Berlin, Germany; Department of Language and Culture, UiT- The Arctic University of Norway, Hansine Hansens veg 18, 9019, Tromsø, Norway; Department of Psychology, Ashoka University, Sonipat, India; India Autism Center, Kolkata, India

**Keywords:** Autism spectrum disorders, autistic traits, grey matter, white matter microstructure, Structural MRI, Diffusion Tensor Imaging

## Abstract

Autism Spectrum Disorders (ASD) are a set of neurodevelopmental conditions characterised by difficulties in social interaction and communication as well as stereotyped and restricted patterns of interest. Autistic traits exist in a continuum across the general population, whilst the extreme end of this distribution is diagnosed as clinical ASD. While many studies have investigated brain structure in autism using a case-control design, few have used a dimensional approach. To add to this growing body of literature, we investigated the structural brain correlates of autistic traits in a mixed sample of adults (N=91) with and without a clinical diagnosis of autism. We examined regional brain volumes (using voxel-based morphometry and surface-based morphometry) and white matter microstructure properties (using Diffusion Tensor Imaging). Our findings show widespread grey matter differences, including in the social brain regions, and some evidence for white matter microstructure differences related to higher autistic traits. These grey matter and white matter microstructure findings from our study are consistent with previous reports and support the brain structural differences in ASD. These findings provide further support for shared aetiology for autistic traits across the diagnostic divide.

## 1. Introduction

Autism Spectrum Disorders (ASD) are complex neurodevelopmental conditions characterised by atypicalities in social interaction and communication as well as stereotyped behaviours (American Psychiatric Association, 2013). The origin of differences in brain structure and volume can be traced back to early childhood, as several studies reported early brain overgrowth in younger children (2-5 years old) with ASD (Courchesne et al., 2001; Hardan et al., 2001). It was suggested that the enlarged brain structural abnormalities were indexed by an increase in head circumference (Courchesne et al., 2003). Such brain structural differences may, in some cases, continue to exist until adulthood in ASD. These differences in brain structure may reflect alternate trajectories of brain development which have consequences for the behavioural manifestations of ASD.

Several studies measuring brain structure in autism have used traditional voxel based morphometry (VBM) (Nickl-Jockschat et al., 2012) and showed reduced regional grey matter volume (GMV) in cortical brain regions including the orbitofrontal cortex (OFC) (Hardan et al., 2006; Mueller et al., 2013), amygdala, fusiform gyrus (FG) (Sato et al., 2017), and superior temporal sulcus (STS) (Boddaert et al., 2004) in individuals with ASD compared to controls (Cauda et al., 2014; Mundy, 2018; Via et al., 2011). These regions are considered to be part of the putative ‘social brain’ circuit and believed to play a significant role in theory of mind abilities, emotional judgement, face recognition and interpreting biological motion cues respectively (Brothers, 1990; Pelphrey et al., 2011; Schultz, 2005). Although regional GMV can be measured using VBM, two-third of the cortical structures are hidden and it may be difficult to directly measure other surface based measures such as cortical thickness, surface area and gyrification by conventional volumetric approaches like VBM (Jiao et al., 2010). Considering this, some previous studies used surface based morphometry (including the regional and inter-regional structural networks) and demonstrated increased cortical thickness in the medial prefrontal cortex and reduced cortical thickness in the posterior cingulate cortex and precuneus in individuals with ASD relative to controls (Valk et al., 2015).

Previous studies have suggested that key brain regions may be poorly connected due to white matter microstructure differences which may affect the social information processing in individuals with ASD (Rippon et al., 2007; Wass, 2011). These differences in the white matter microstructure may be driven by reduced axonal density and myelination in individuals with ASD. Previous studies on ASD have reported white matter microstructure abnormalities (reduced fractional anisotropy and increased mean diffusivity) in the fibre tracts including bilateral superior longitudinal fasciculus (SLF), uncinate fasciculus (UF), inferior longitudinal fasciculus (ILF) and inferior fronto-occipital fasciculus (IFOF) (Boets et al., 2018; Catani et al., 2016; Itahashi et al., 2015; Barnea-Goraly et al., 2004; Groen et al., 2011; Lee et al., 2007; Lisiecka et al., 2015) which connects key social brain regions. These differences in the regional grey matter (including the social brain regions) and white matter microstructure (Aoki et al., 2013) indicate brain structural atypicalities are believed to play an important role in individuals with ASD.

The majority of the studies discussed above have used a case-control design and reported brain structural differences including regional grey matter volume (GMV) and white matter microstructure in individuals with ASD. These brain structure measurements between the ASD and control group may induce a sampling bias in the analysis and lead to mixed findings. There is considerable variance inherent in the case-control design due to the sampling of the controls. A dimensional approach avoids this source of variance by sampling across the whole population. Growing evidence suggests that autistic traits lie in a continuum across the general population, whilst the higher end of this trait measure is diagnosed as clinical ASD (Whitehouse et al., 2011; Robinson et al., 2011). More importantly, it is very important to understand the brain structure, rather than solely depending upon the behavioural measures assessing the symptoms of autism irrespective of categorising clinical ASD. This is because although ASD is a behaviourally defined condition, from the previous studies, brain structural differences are believed to underlie the atypical behavioural manifestations in ASD. Taking all these factors into account, we used surface-based morphometry to measure the cortical thickness, surface area and gyrification in the cortical grey matter, and we used voxel based morphometry to characterise both cortical and subcortical grey matter volume at a whole brain level. In addition, we also examined the white matter microstructure differences using DTI. We explored the relationship of these brain based metrics with self-reported autistic traits in adults with and without a diagnosis of ASD.

## 2. Methods and materials

### 2.1 Participants

Ninety-one adults consisting of 66 neurotypicals and 25 ASD (52 males, 39 females, age 18-60 years), participated in this study. All neurotypical individuals were recruited from the University of Reading campus and individuals with ASD were recruited from a registered clinic. Participants with symptoms of ASD were assessed using the Autism Diagnostic Observation Schedule (ADOS) module-4 and diagnosed using the diagnostic and statistical manual (DSM IV-TR). Subjects with any neurological conditions or head injuries were excluded from the study. Autism Spectrum Quotient (AQ) scores (Baron-Cohen et al., 2001) were also collected from all participants. This dataset came from two separate phases of data collection (N=53 and N= 38; both phases used the same protocol for collecting structural MRI). This study was approved by the University Research Ethics Committee (UREC), University of Reading. From the full sample above, a subset of fifty-three adults consisting 28 neurotypicals and 25 ASD (31 males and 22 females, age 18-60 years) matched for age, gender and IQ took part in the diffusion tensor imaging study. The performance IQ was measured using Raven’s Progressive Matrices (RPM) (Raven, 2000) to ensure including participants with no intellectual disabilities.

### 2.2 sMRI and DTI Data collection

Siemens Trio 3T MRI Scanner was used to acquire the high resolution T1-weighted whole brain structural images from all participants using 32-channel head coil including (Voxel size = 1 × 1 × 1 mm^3^; matrix =256 × 256; TR = 2020ms; TE = 2ms) at the Centre for Integrative Neurosciences and Neurodynamics (CINN). The DTI protocol used single-shot spin echo, echo planar imaging (EPI) with 32-gradients including 60 diffusion weighted (b=1000 sec/mm^2^) and 2 non-diffusion weighted images (b=0 sec/mm2), repetition time = 7200 ms; echo time = 10 ms, matrix =128 × 128, voxel size = 2 × 2 × 2 mm (isotropic).

### 2.3 SBM preprocessing

Freesurfer analysis suite (Fischl, 2012) was used to perform the surface-based morphometry to reconstruct the cortical surface. The MPRAGE images were preprocessed and corrected for head motion, bias field correction, skull-stripping, segmentation, registration, spatial normalisation and smoothing. After the bias-field correction and skull-stripping, the individual structural images were computed to determine the transformation matrix and co-registered to the Talairach space to maximise the possibility that individual images overlap with the study-specific average brain template coordinates. Then, the structural brain images were segmented into pial and white surfaces. Next, the inflated cortical surfaces from the individual images were spatially normalised to the spherical average template, such that each vertex forming multiple triangles across the surface were aligned closely to the corresponding anatomical locations. In the final step, default smoothing was applied to normalise the local neighbourhood voxels across the entire brain. The pial surfaces of each hemisphere were preprocessed to create an outer smoothed pial surface to account for the local gyrification index (LGI).

All the individual subjects’ cortical thickness, surface area and local gyrification index maps were concatenated together for measuring each metric separately in the group level analysis. Additionally, smoothing (FWHM= 10mm) was applied to average the close neighbourhood voxels for cortical thickness, surface area, while no additional smoothing was used for measuring the local gyrification index in this analysis based on its compatibility of the cluster-forming threshold (0.05).

### 2.4 VBM preprocessing

Voxel-based morphometry was applied for preprocessing and analysis with the Diffeomorphic Anatomical Registration Through Exponentiated Lie algebra (DARTEL) pipeline incorporated in SPM12 toolbox. Initially the T1-weighted structural brain (MPRAGE) images were reoriented for anterior and posterior commissure alignment and corrected for head motion. In this method, images were segmented into grey matter, white matter and CSF. Then, a study-specific template was created by aligning and averaging the inter-subject grey matter volumes iteratively. The segmented individual grey matter volumes were registered to the template using non-linear registration and normalised to MNI standard space. These normalised images were smoothed using Gaussian kernel (FWHM=8mm) for the cortical structures and subcortical structures (FWHM=4mm) by averaging the spatial intensity of the local neighbouring voxels (Ashburner, 2010; Coalson et al., 2018).

### 2.5 Tract-based Spatial Statistics

Tract-based Spatial Statistics (TBSS), a whole-brain voxel-wise analytical approach incorporated in the FSL software library version 5.0 (Smith et al., 2006) was used for the data analysis. The standard TBSS preprocessing and analysis pipeline was used for eddy current correction, non-brain tissue removal, diffusion tensor modelling, registration, normalisation, thresholding and randomisation as follows: The DTI images were preprocessed for eddy current correction and removal of non-brain tissues, and the diffusion tensor models (FA and MD maps) were derived from all the images. Subsequently, the FA and MD maps were non-linearly registered and transformed to the FA FMRIB (1mm^3^) standard space. Next, the mean FA skeleton, all skeletonised FA and MD 4D concatenated multi-subject maps were derived and transformed using non-linear registration to the MNI152 (1mm^3^) standard space. Then, a white matter thresholding (0.2) was used on the mean FA skeleton to restrict the grey matter partial volume effects.

## 3. Statistical analysis

### SBM analysis

In the SBM analysis, the Different Offset Same Slope model was used to test the relationship between cortical thickness, surface area, local gyrification index (separately for each dependent variable at a time) and AQ, including age and gender as covariates. The FreeSurfer Group Descriptor format was used to construct a design matrix. Then, the precomputed Monte-Carlo Simulation was used to run the tests for multiple comparisons with a cluster-forming threshold (0.05) and the threshold for significance (p=0.05, two-tailed).

### VBM analysis

The general linear model was used to test the relationship between the regional GMV and AQ scores across the combined sample of ASD and neurotypicals after controlling for the effects of age, gender and total brain volume. The covariates including the age and gender were demeaned for the whole sample. We used Family Wise Error (FWE) rate testing for multiple comparisons.

### Tract-based spatial statistics

The general linear model was used to test the relationship between fractional anisotropy and AQ. In addition, the relationship between the mean diffusivity and AQ was also tested, while controlling for age, gender and IQ. This was performed using permutation-based testing (N=5000) and Threshold-free Cluster Enhancement (TFCE) for multiple comparisons.

## 4. Results

### Surface based morphometry

Our analysis focused on the relationship between surface-based morphometry and autistic traits revealed significant positive association between all four metrics including cortical thickness, surface area, cortical volume, local gyrification index and autistic traits across the combined sample of neurotypicals and individuals with ASD. Autistic traits were found to be significantly associated with cortical thickness in the left lingual gyrus, right lateral occipital cortex and right pars triangularis, and with surface area in the right lateral occipital cortex. The associations with cortical volume were observed in the left lingual gyrus, right lateral occipital cortex and right pars triangularis. In addition, the significantly associated clusters for local gyrification index were observed in the right lingual gyrus (Table 2).

**Table 1:**
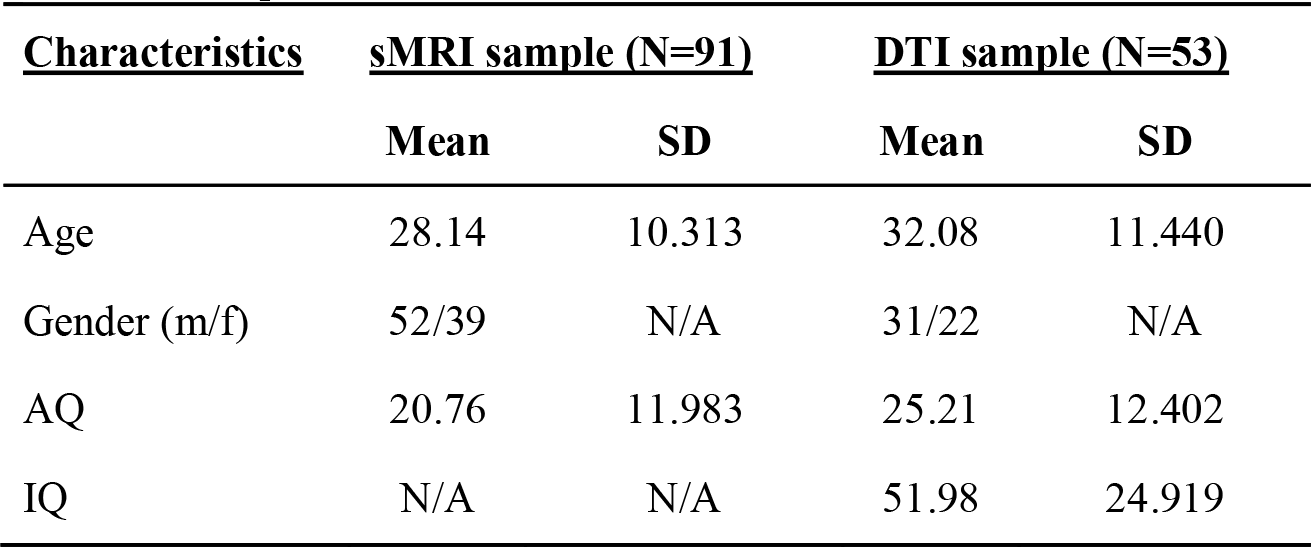
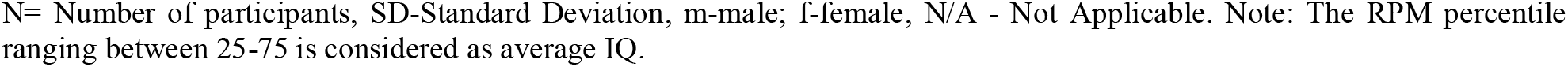
Sample characteristics

| **<u>Characteristics</u>** | **<u>sMRI sample (N=91)</u>** |  | **<u>DTI sample (N=53)</u>** |  |
| --- | --- | --- | --- | --- |
|  | **Mean** | **SD** | **Mean** | **SD** |
| Age | 28.14 | 10.313 | 32.08 | 11.440 |
| Gender (m/f) | 52/39 | N/A | 31/22 | N/A |
| AQ | 20.76 | 11.983 | 25.21 | 12.402 |
| IQ | N/A | N/A | 51.98 | 24.919 |

N= Number of participants, SD-Standard Deviation, m-male; f-female, N/A - Not Applicable. Note: The RPM percentile ranging between 25-75 is considered as average IQ.

**Table 2:** Brain showing association between cortical thickness, surface area, cortical volume and AQ

| <u>Brain regions</u> | <u>Hem</u> | <u>Talairach coordinates</u> |  |  | <u>t-value</u> | <u>K</u> |
| --- | --- | --- | --- | --- | --- | --- |
|  |  | x | y | z |  |  |
| Cortical thickness |  |  |  |  |  |  |
| Lingual gyrus | L | -14.2 | -63.4 | -3.5 | 3.642 | 2386.74 |
| Lateral Occipital | R | 18.3 | -100.3 | -7 | 4.255 | 4388.31 |
| Pars triangularis | R | 47 | 35 | 4.7 | 3.679 | 1480.64 |
| Surface area |  |  |  |  |  |  |
| Lateral Occipital | R | 16.2 | -99.1 | -2.5 | 3.434 | 4042.26 |
| Local gyrification index |  |  |  |  |  |  |
| Lingual gyrus | R | 18.6 | -64.2 | -8.3 | 2.493 | 2392.50 |

Abbreviations: L- Left, R- Right, K-cluster size, t-value - test statistics, Hem-Hemisphere, p-threshold = 0.05, corrected.

**Fig. 1:**
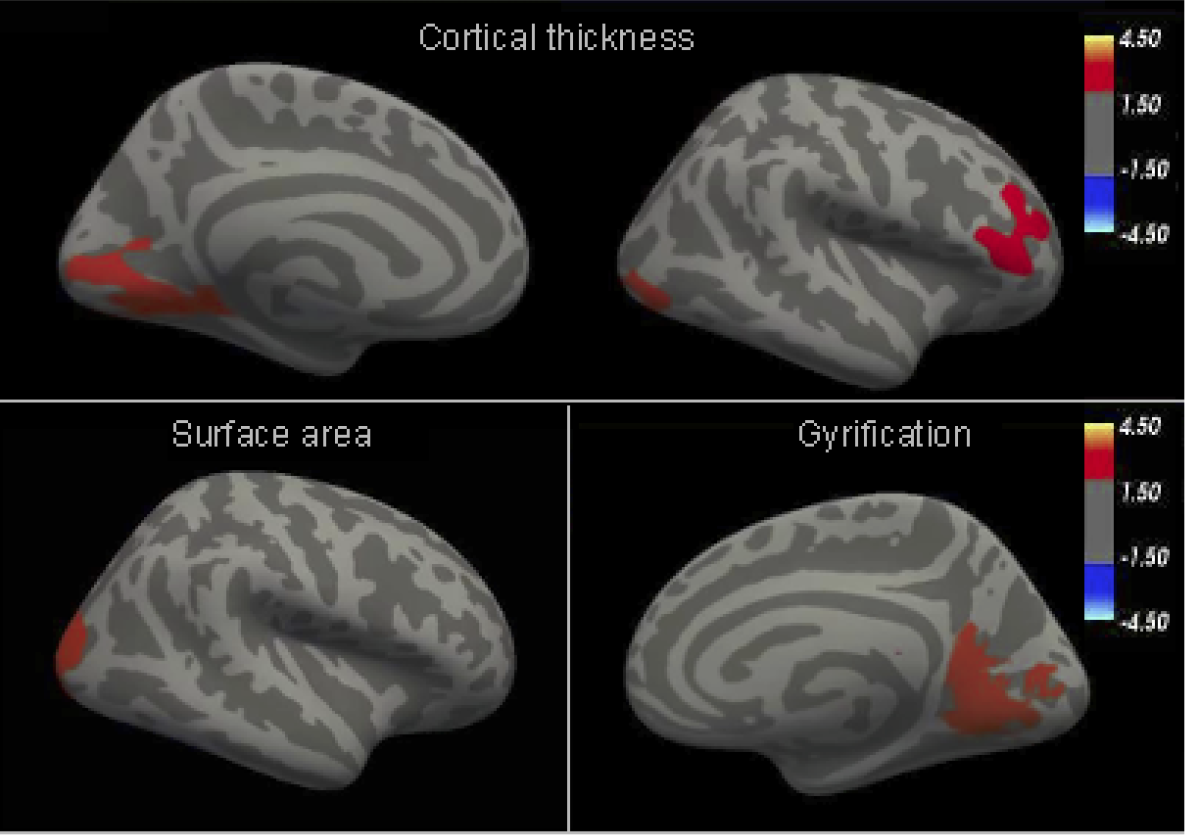
**Cortical thickness:** Clusters showing significantly associated brain regions of lingual Gyrus (left), lateral occipital (Right) and pars triangularis (right). Cortical thickness is measured in millimetres (mm). **Surface area:** Clusters showing significantly associated brain regions in lateral occipital cortex (right). The unit of surface area is square millimetre (mm^2^). **LGI:** Clusters showing significantly associated brain regions in lingual gyrus (right). Local gyrification index has no units.

**Fig. 2:**
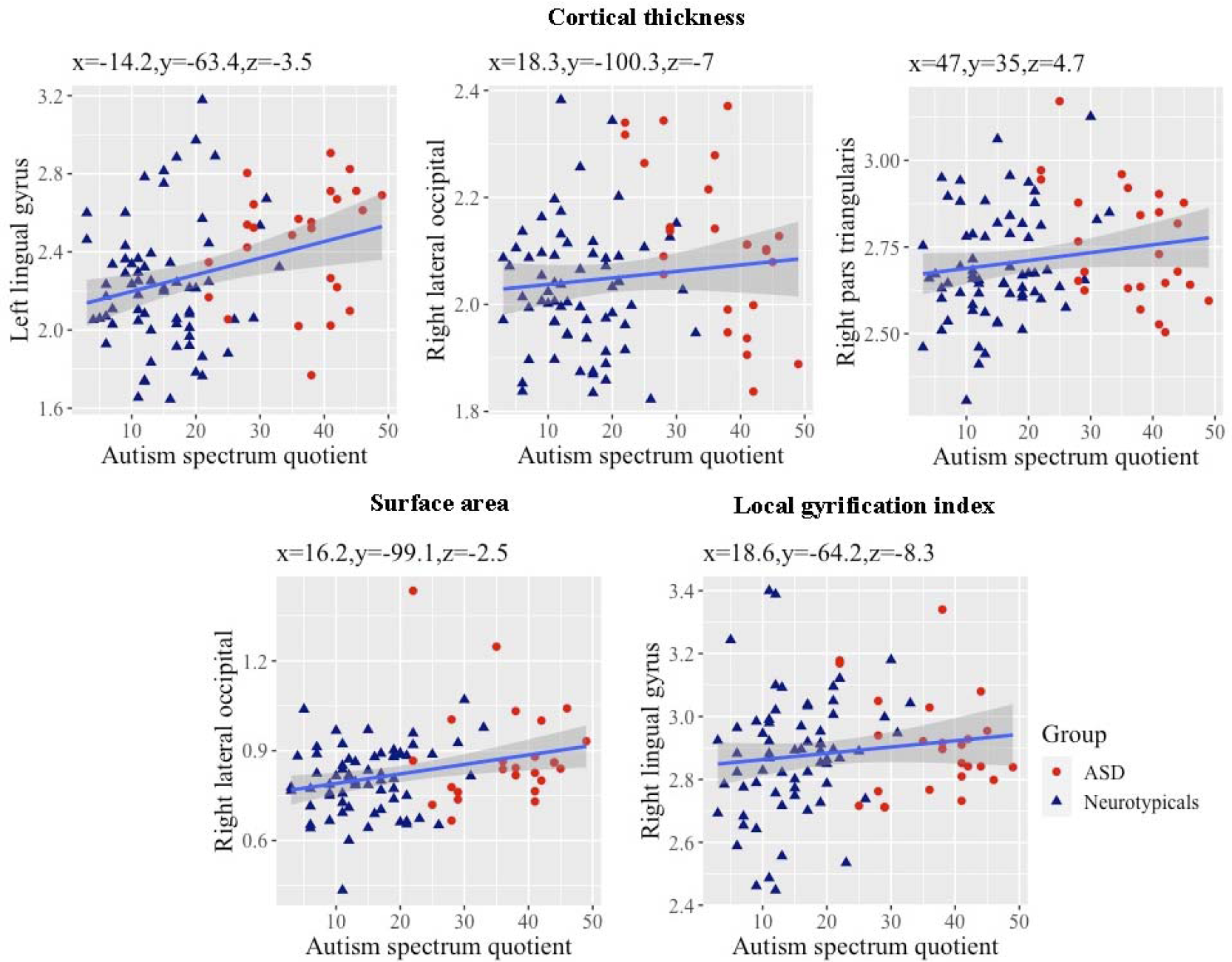
Scatterplot showing positive association between cortical thickness (first row), surface area (second row, left), gyrification (second row, right) in different brain regions with AQ scores. The colored triangles in navy blue and red indicate controls and ASD respectively.

### Voxel based morphometry

We found significant positive association between regional GMV and AQ scores in cortical brain regions including the clusters of right lingual gyrus and precentral gyrus. We also found significant positive association between regional GMV and AQ scores in subcortical brain regions including the left putamen and right putamen. Additionally, we found significant negative association between regional GMV and AQ in the right orbitofrontal cortex which also extended to the anterior cingulate gyrus. (Table 3).

**Table 3:** Brain regions displaying the association between cortical and subcortical GMV and AQ

| Cortical brain regions | Hem | MNI-Coordinates |  |  | K | P<sub>FWE</sub> |
| --- | --- | --- | --- | --- | --- | --- |
|  |  | x | y | z |  |  |
| Positive association |  |  |  |  |  |  |
| Lingual gyrus | R | 9 | -65 | -8 | 8010 | <0.001*** |
| Precentral gyrus | R | 14 | -22 | 62 | 6918 | 0.012** |
| Negative association |  |  |  |  |  |  |
| Orbitofrontal cortex | R | 16 | 21 | -25 | 7888 | <0.001*** |
| Subcortical brain regions |  |  |  |  |  |  |
| Positive association |  |  |  |  |  |  |
| Putamen | L | -29 | -9 | -3 | 2837 | <0.001*** |
| Putamen | R | 29 | -4 | -5 | 2459 | <0.001*** |

Abbreviations: Level of significance \*p < .05, \*\*p < .01, \*\*\*p < .001, Hem - Hemisphere, FWE- Family Wise Error and K- Cluster size.

**Fig. 3:**
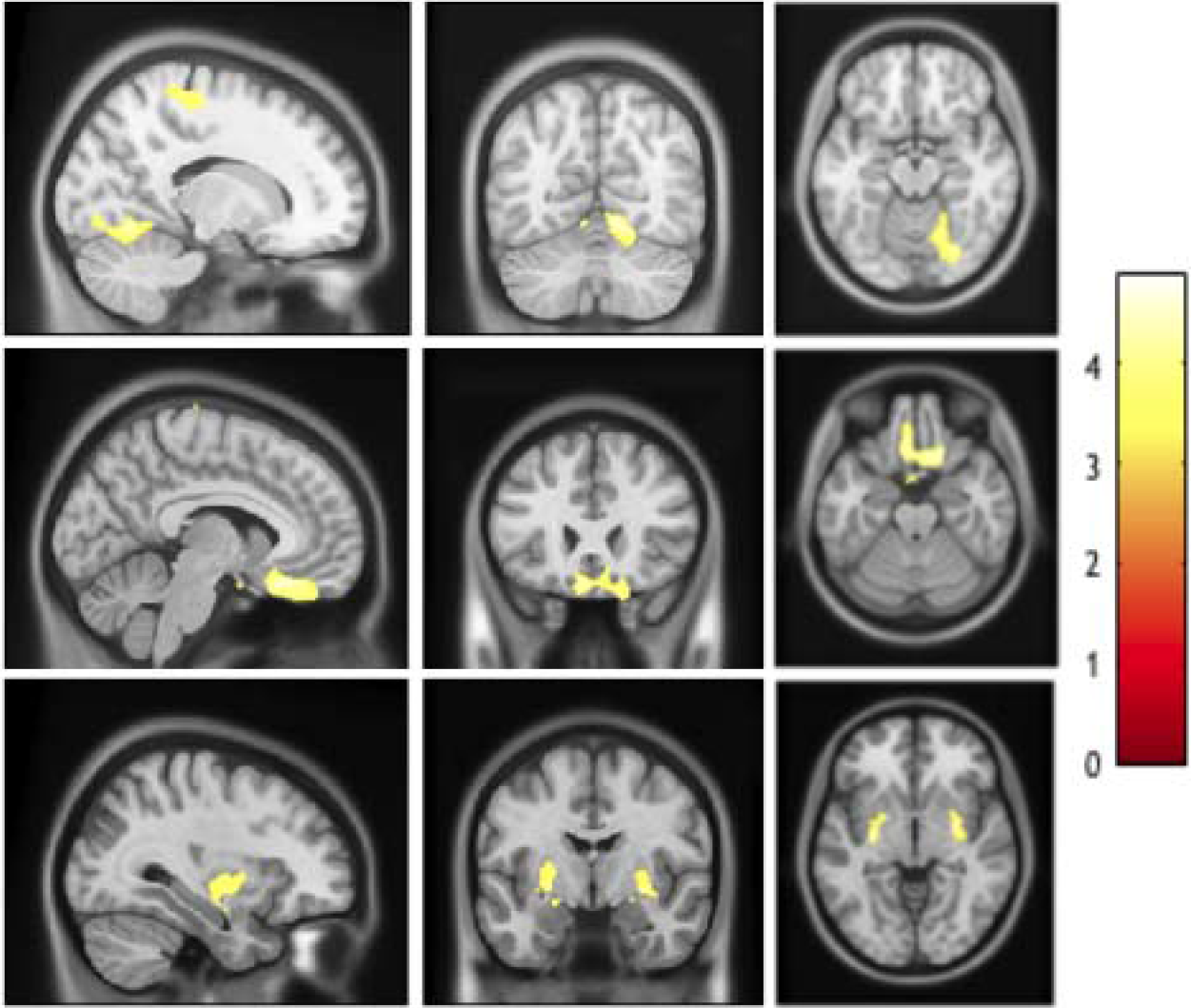
Row 1: Significantly associated cluster in right lingual gyrus and right precentral gyrus. Row 2: Significantly associated cluster in right orbitofrontal cortex. Row 3: Significantly associated clusters in bilateral putamen.

**Fig. 4:**
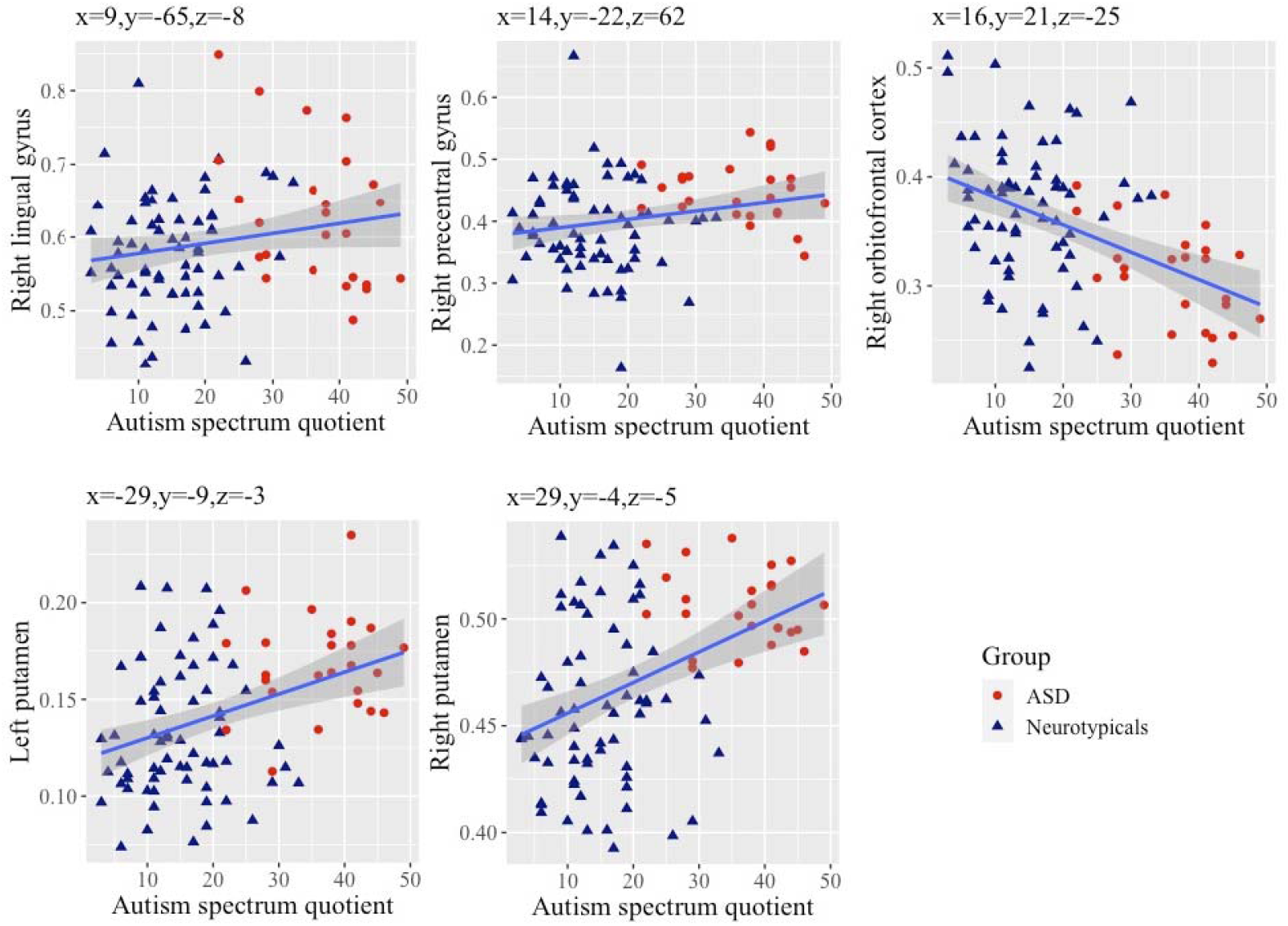
Scatterplots showing significant positive association between GMV of different brain regions including right lingual gyrus, right precentral gyrus, left and right putamen with AQ scores. Scatterplot showing significant negative association between right orbitofrontal cortex GMV and AQ scores.

### Tract based Spatial Statistics

We found a positive association between MD and AQ in the superior longitudinal fasciculus, inferior longitudinal fasciculus, inferior fronto-occipital fasciculus and corpus callosum (forceps major and splenium). In addition, we also found a negative association between FA and AQ in the superior longitudinal fasciculus, inferior longitudinal fasciculus, inferior fronto-occipital fasciculus and corticospinal tract in the combined sample of neurotypicals and individuals with ASD. However, none of these clusters survived after correcting for multiple comparisons using threshold-free cluster enhancement (TFCE) (p < 0.05, uncorrected) (Table 4).

**Table 4:**
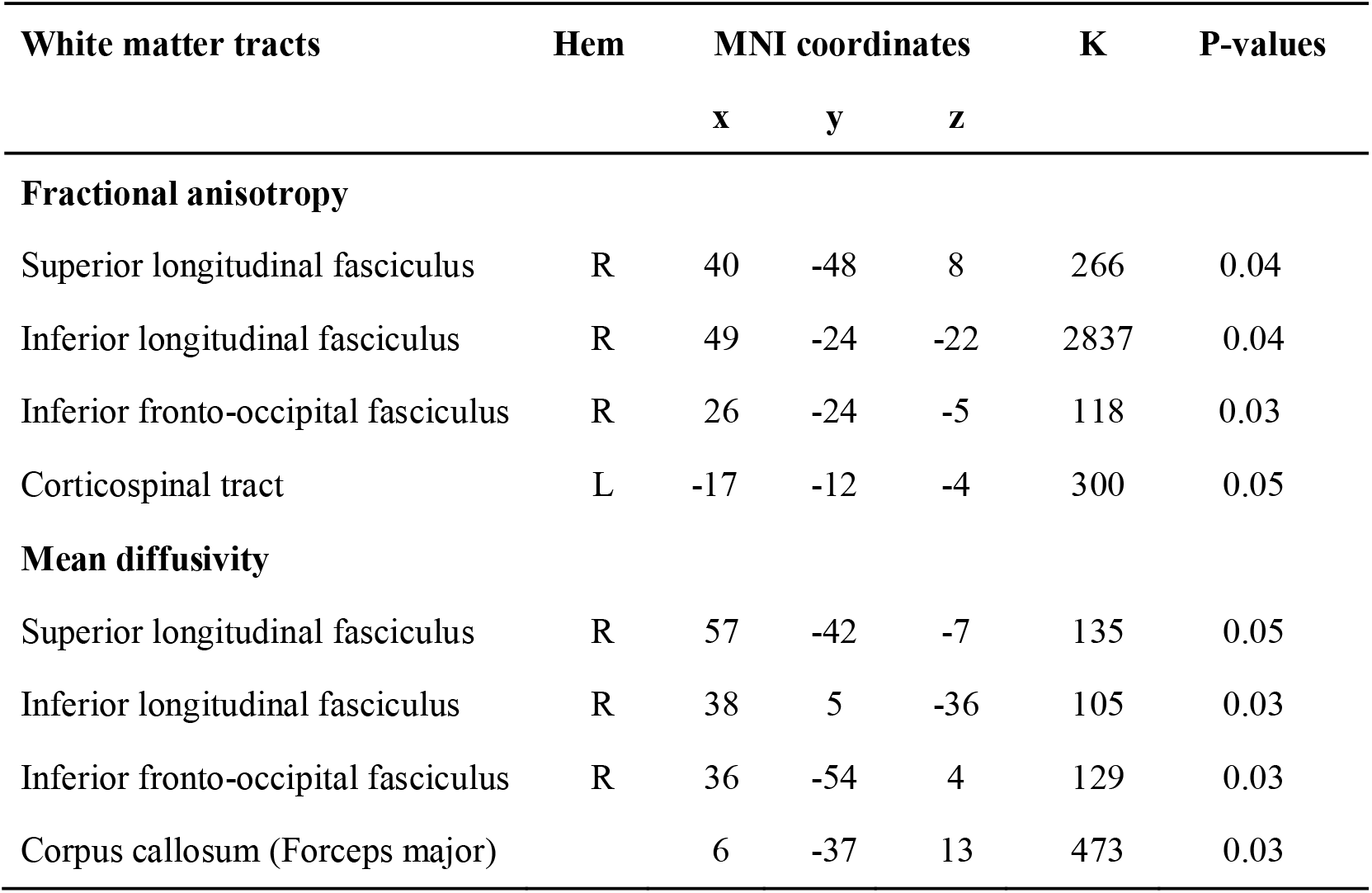
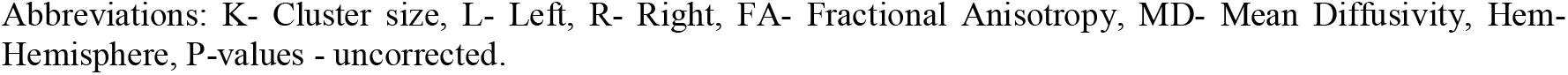
Association between fractional anisotropy and mean diffusivity and AQ

| White matter tracts | Hem | MNI coordinates |  |  | K | P-values |
| --- | --- | --- | --- | --- | --- | --- |
|  |  | x | y | z |  |  |
| Fractional anisotropy |  |  |  |  |  |  |
| Superior longitudinal fasciculus | R | 40 | -48 | 8 | 266 | 0.04 |
| Inferior longitudinal fasciculus | R | 49 | -24 | -22 | 2837 | 0.04 |
| Inferior fronto-occipital fasciculus | R | 26 | -24 | -5 | 118 | 0.03 |
| Corticospinal tract | L | -17 | -12 | -4 | 300 | 0.05 |
| Mean diffusivity |  |  |  |  |  |  |
| Superior longitudinal fasciculus | R | 57 | -42 | -7 | 135 | 0.05 |
| Inferior longitudinal fasciculus | R | 38 | 5 | -36 | 105 | 0.03 |
| Inferior fronto-occipital fasciculus | R | 36 | -54 | 4 | 129 | 0.03 |
| Corpus callosum (Forceps major) |  | 6 | -37 | 13 | 473 | 0.03 |

Abbreviations: K- Cluster size, L- Left, R- Right, FA- Fractional Anisotropy, MD- Mean Diffusivity, Hem- Hemisphere, P-values - uncorrected.

**Fig. 5:**
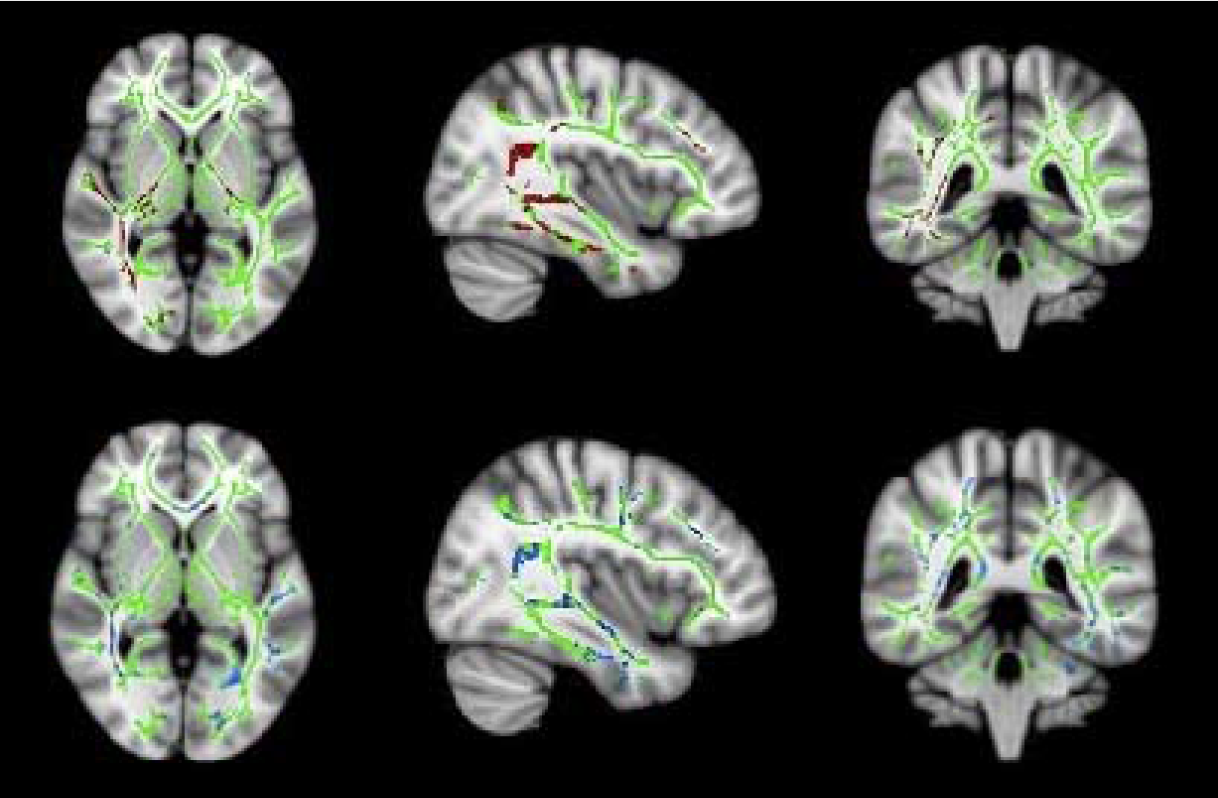
Whole white matter skeleton (green) display, showing negative association between fractional anisotropy (clusters in red, first row) with AQ and positive association between mean diffusivity (clusters in blue, second row) with AQ (4).

## 5. Discussion

In the current study, we tested the relationship between the regional grey matter properties, white matter microstructure and autistic traits in a mixed sample with adults with and without a clinical diagnosis of ASD. Our results demonstrated that autistic traits were significantly associated with multiple metrics of regional grey matter (cortical thickness, surface area, gyrification and volume) that spanned the social brain regions. These findings were consistent with brain structural findings from previous studies that used a case-control design (Ecker et al., 2012; Sato et al., 2017; Shukla et al., 2011).

We discovered significant regional grey matter variations in the key social brain regions in the frontal lobe related to higher autistic traits. These findings include reduced regional GMV in the right orbitofrontal cortex and increased cortical thickness in the right pars triangularis. The reduced regional GMV in the orbitofrontal cortex may be underpinned by fewer minicolumns in the frontal lobes demonstrated in post-mortem studies (Buxhoeveden et al., 2006; Casanova et al., 2006). The regional GMV differences in the orbitofrontal cortex has previously been suggested to be related to observed behavioural differences in theory of mind (ToM) in individuals with ASD (Frith & Frith, 2001; Lewis et al., 2011; Sabbagh, 2004). Individuals with higher autistic traits might have social skill difficulties such as interpreting self-thoughts and interpreting other’s intentions (Girgis et al., 2007; Mundy, 2003). The greater cortical thickness in the pars triangularis may be related to the expressive language deficits noted in some individuals with ASD (Knaus et al., 2018). The pars triangularis may cross-talk with the other social brain region (pars orbitalis) closely located in the frontal lobe, which may also account for the social communication difficulties related to higher autistic traits (Fishman et al., 2014).

Greater GMV in the precentral gyrus (in the right hemisphere) was associated with higher autistic traits. This finding is consistent with previous studies in ASD relative to controls (Bonilha et al., 2008; Ecker et al., 2012; Rojas et al., 2006). The precentral gyrus is believed to be an integral part of an action observation network/mirror neuron system (Hadjikhani et al., 2006). Increased GMV in the precentral gyrus may underlie atypical visuomotor learning in individuals with higher autistic traits (Mahajan et al., 2016; Carper & Courchesne, 2005; Nebel et al., 2014). We also found increased regional GMV in the bilateral putamen related to higher autistic traits. This finding is consistent with regional GMV variations in putamen in ASD from previous studies (Hollander et al., 2005; Langen et al., 2009; Nickl-Jockschat et al., 2012). The putamen, an integral part of the dorsal striatum, plays a key role in restricted and repetitive behaviour in ASD (Langen et al., 2012; Sato et al., 2014). Such regional GMV variations in putamen may influence the striatum volume which may underlie atypical behavioural manifestations such as insistence to sameness and complex motor functions in individuals with ASD (Calderoni et al., 2014; Eisenberg et al., 2015; Schuetze et al., 2016). These common sites of brain structural variations in the striatum may underlie the stereotyped behaviours in individuals with ASD (Eisenberg et al., 2015; Schuetze et al., 2016) which may also be related to higher autistic traits.

The lingual gyrus in the right hemisphere demonstrated increased gyrification and regional grey matter volume related to higher autistic traits. In addition, lingual gyrus in the left hemisphere demonstrated increased cortical thickness. This evidence is consistent with previous reports of structural atypicalities, with greater local gyrification and grey matter volume in lingual gyrus in individuals with ASD (Libero et al., 2019; Peterson et al., 2006). In addition, the right lateral occipital cortex showed cortical thickness and increased surface area which may result in difficulties when modulating the visual perceptual abilities (Ecker et al., 2010). Lingual gyrus constitutes part of a network, including other brain regions (lateral occipital cortex, fusiform gyrus and posterior superior temporal sulcus) that play a significant role in object/face recognition and following biological motion cues in ASD (Ecker et al., 2015). The lateral occipital cortex is believed to play a significant role in visuospatial attention in individuals with ASD (Ecker et al., 2013; Nickl-Jockschat et al., 2012). The greater volume and gyrification of the lingual gyrus and lateral occipital cortex may underlie the atypical visual processing in individuals with higher autistic symptoms (Keehn et al., 2008).

The regional variations in intrinsic grey matter properties may arise from differences in neuronal migration within the radial minicolumns which may be altered in individuals with ASD/higher autistic traits (Casanova & Trippe, 2009). This aberrant cortical cytoarchitecture may be indexed by an increased number of minicolumns, reduced alignment and increased density of pyramidal neuronal cells - and may be a key factor associated with the atypical cortico-cortical connectivity in ASD. These developmental neurobiological processes may underlie the observed pattern of brain structural metrics in the pars triangularis, lateral occipital and lingual gyrus that are associated with higher autistic traits. Our findings from VBM and SBM study supports the evidence for variations in regional brain volume and atypical cortico-cortical connectivity hypothesis in ASD.

Notably, there are some methodological differences between VBM and SBM (topographical and voxel-wise comparison respectively) in measuring cortical morphometry (Hyde et al., 2010; Jiao et al., 2010; Pappaianni et al., 2018). These two analytical approaches (VBM and SBM) are incomparable when measuring the cortical thickness, surface area and gyrification because their principles and implementation are distinct from one another. SBM provides us with a higher reliability in measuring the cortical thickness, surface and gyrification, whereas the VBM (DARTEL) provides us with a high dimensional spatial registration for measuring regional grey matter volume in ASD. In addition, VBM helps us to measure the regional GMV in the subcortical structures unlike SBM. Together, VBM and SBM are the two complementary approaches that contribute to the efforts in identifying a neuroimaging endophenotype for ASD.

While none of the DTI results survived a test for multiple comparisons, the findings from the DTI study were convergent with those from the SBM study. The SBM study found atypicalities in the lingual gyrus, which is connected to the ventral visual stream through the ILF and IFOF. These findings suggest that the co-occurrence of grey matter variations of these brain regions (lingual gyrus and lateral occipital cortex) and atypical white matter microstructure integrity of inferior longitudinal fasciculus (ILF) and inferior fronto-occipital fasciculus (IFOF) may underlie the sensory atypicalities in individuals with higher autistic traits (Itahashi et al., 2015). In addition, the white matter microstructure variations in the superior longitudinal fasciculus (SLF) (connected to the pars triangularis and wernicke’s area) may impose difficulties in acquiring language skills may be associated with autistic symptoms (Fitzgerald et al., 2018).

## 6. Conclusion

The regional grey matter variations in the orbitofrontal cortex and pars triangularis, dorsal striatum and ventral visual stream were found to be related to higher autistic traits. These observations are consistent with previous results reported in case-control studies of ASD. Our study provides further evidence in support of a dimensional approach to investigate potential neural endophenotypes related to autism.

## References

American Psychiatric Association. (2013). Diagnostic and Statistical Manual of Mental Disorders (DSM-5®). American Psychiatric Pub. https://play.google.com/store/books/details?id=-JivBAAAQBAJ

Aoki, Y., Abe, O., Nippashi, Y., & Yamasue, H. (2013). Comparison of white matter integrity between autism spectrum disorder subjects and typically developing individuals: a meta-analysis of diffusion tensor imaging tractography studies. Molecular autism, 4(1), 1–17. https://doi.org/10.1186/2040-2392-4-25

Ashburner, J. (2010). VBM tutorial. Tech. repWellcome Trust Centre for Neuroimaging, London, UK. https://www.fil.ion.ucl.ac.uk/~john/misc/VBMclass15.pdf

Barnea-Goraly, N., Kwon, H., Menon, V., Eliez, S., Lotspeich, L., & Reiss, A. L. (2004). White matter structure in autism: preliminary evidence from diffusion tensor imaging. In Biological Psychiatry (Vol. 55, Issue 3, pp. 323–326). https://doi.org/10.1016/j.biopsych.2003.10.022

Baron-Cohen, S., Wheelwright, S., Skinner, R., Martin, J., & Clubley, E. (2001). The autism-spectrum quotient (AQ): Evidence from asperger syndrome/high-functioning autism, males and females, scientists and mathematicians. Journal of Autism and Developmental Disorders, 31(1), 5–17.

Bedford, S. A., Park, M. T. M., Devenyi, G. A., Tullo, S., Germann, J., Patel, R., Anagnostou, E., Baron-Cohen, S., Bullmore, E. T., Chura, L. R., Craig, M. C., Ecker, C., Floris, D. L., Holt, R. J., Lenroot, R., Lerch, J. P., Lombardo, M. V., Murphy, D. G. M., Raznahan, A., … MRC AIMS Consortium. (2020). Large-scale analyses of the relationship between sex, age and intelligence quotient heterogeneity and cortical morphometry in autism spectrum disorder. Molecular Psychiatry, 25(3), 614–628. https://doi.org/10.1038/s41380-019-0420-6

Boddaert, N., Chabane, N., Gervais, H., Good, C. D., Bourgeois, M., Plumet, M.-H., Barthélémy, C., Mouren, M.-C., Artiges, E., Samson, Y., Brunelle, F., Frackowiak, R. S. J., & Zilbovicius, M. (2004). Superior temporal sulcus anatomical abnormalities in childhood autism: a voxel-based morphometry MRI study. NeuroImage, 23(1), 364–369. https://doi.org/10.1016/j.neuroimage.2004.06.016

Boets, B., Van Eylen, L., Sitek, K., Moors, P., Noens, I., Steyaert, J., Sunaert, S., & Wagemans, J. (2018). Alterations in the inferior longitudinal fasciculus in autism and associations with visual processing: a diffusion-weighted MRI study. Molecular Autism, 9, 10. https://doi.org/10.1186/s13229-018-0188-6

Bonilha, L., Cendes, F., Rorden, C., Eckert, M., Dalgalarrondo, P., Li, L. M., & Steiner, C. E. (2008). Gray and white matter imbalance--typical structural abnormality underlying classic autism? Brain & Development, 30(6), 396–401. https://doi.org/10.1016/j.braindev.2007.11.006

Brothers, & L. (1990). The social brain : A project for integrating primate behaviour and neurophysiology in a new domain. Concepts Neurosci, 1, 27–51. https://ci.nii.ac.jp/naid/20000695363/

Buxhoeveden, D. P., Semendeferi, K., Buckwalter, J., Schenker, N., Switzer, R., & Courchesne, E. (2006). Reduced minicolumns in the frontal cortex of patients with autism. Neuropathology and Applied Neurobiology, 32(5), 483–491. https://doi.org/10.1111/j.1365-2990.2006.00745.x

Calderoni, S., Bellani, M., Hardan, A. Y., Muratori, F., & Brambilla, P. (2014). Basal ganglia and restricted and repetitive behaviours in Autism Spectrum Disorders: current status and future perspectives. Epidemiology and Psychiatric Sciences, 23(3), 235–238. https://doi.org/10.1017/S2045796014000171

Carper, R. A., & Courchesne, E. (2005). Localized enlargement of the frontal cortex in early autism. Biological Psychiatry, 57(2), 126–133. https://doi.org/10.1016/j.biopsych.2004.11.005

Casanova, M., & Trippe, J. (2009). Radial cytoarchitecture and patterns of cortical connectivity in autism. Philosophical Transactions of the Royal Society of London. Series B, Biological Sciences, 364(1522), 1433–1436. https://doi.org/10.1098/rstb.2008.0331

Casanova, M. F., van Kooten, I. A., Switala, A. E., van Engeland, H., Heinsen, H., Steinbusch, H. W., … & Schmitz, C. (2006). Minicolumnar abnormalities in autism. Acta neuropathologica, 112(3), 287. DOI 10.1007/s00401-006-0085-5

Catani, M., Dell’Acqua, F., Budisavljevic, S., Howells, H., Thiebaut de Schotten, M., Froudist-Walsh, S., D’Anna, L., Thompson, A., Sandrone, S., Bullmore, E. T., Suckling, J., Baron-Cohen, S., Lombardo, M. V., Wheelwright, S. J., Chakrabarti, B., Lai, M.-C., Ruigrok, A. N. V., Leemans, A., Ecker, C., … Murphy, D. G. M. (2016). Frontal networks in adults with autism spectrum disorder. Brain: A Journal of Neurology, 139(Pt 2), 616–630. https://doi.org/10.1093/brain/awv351

Cauda, F., Costa, T., Palermo, S., D’Agata, F., Diano, M., Bianco, F., Duca, S., & Keller, R. (2014). Concordance of white matter and gray matter abnormalities in autism spectrum disorders: a voxel-based meta-analysis study: Concordance of WM and GM Abnormalities in ASD. Human Brain Mapping, 35(5), 2073–2098. https://doi.org/10.1002/hbm.22313

Coalson, T. S., Van Essen, D. C., & Glasser, M. F. (2018). The impact of traditional neuroimaging methods on the spatial localization of cortical areas. Proceedings of the National Academy of Sciences of the United States of America, 115(27), E6356–E6365. https://doi.org/10.1073/pnas.1801582115

Courchesne, E., Carper, R., & Akshoomoff, N. (2003). Evidence of brain overgrowth in the first year of life in autism. JAMA: The Journal of the American Medical Association, 290(3), 337–344. https://doi.org/10.1001/jama.290.3.337

Courchesne, E., Karns, C. M., Davis, H. R., Ziccardi, R., Carper, R. A., Tigue, Z. D., Chisum, H. J., Moses, P., Pierce, K., Lord, C., Lincoln, A. J., Pizzo, S., Schreibman, L., Haas, R. H., Akshoomoff, N. A., & Courchesne, R. Y. (2001). Unusual brain growth patterns in early life in patients with autistic disorder: an MRI study. Neurology, 57(2), 245–254. https://doi.org/10.1212/wnl.57.2.245

Ecker, C., Bookheimer, S. Y., & Murphy, D. G. (2015). Neuroimaging in autism spectrum disorder: brain structure and function across the lifespan. The Lancet Neurology, 14(11), 1121–1134. https://doi.org/10.1016/S1474-4422(15)00050-2

Ecker, C., Ginestet, C., Feng, Y., Johnston, P., Lombardo, M. V., Lai, M.-C., Suckling, J., Palaniyappan, L., Daly, E., Murphy, C. M., Williams, S. C., Bullmore, E. T., Baron-Cohen, S., Brammer, M., Murphy, D. G. M., & MRC AIMS Consortium. (2013). Brain surface anatomy in adults with autism: the relationship between surface area, cortical thickness, and autistic symptoms. JAMA Psychiatry, 70(1), 59–70. https://doi.org/10.1001/jamapsychiatry.2013.265

Ecker, C., Rocha-Rego, V., Johnston, P., Mourao-Miranda, J., Marquand, A., Daly, E. M., Brammer, M. J., Murphy, C., Murphy, D. G., & MRC AIMS Consortium. (2010). Investigating the predictive value of whole-brain structural MR scans in autism: a pattern classification approach. NeuroImage, 49(1), 44–56. https://doi.org/10.1016/j.neuroimage.2009.08.024

Ecker, C., Suckling, J., Deoni, S. C., Lombardo, M. V., Bullmore, E. T., Baron-Cohen, S., Catani, M., Jezzard, P., Barnes, A., Bailey, A. J., Williams, S. C., Murphy, D. G. M., & MRC AIMS Consortium. (2012). Brain anatomy and its relationship to behavior in adults with autism spectrum disorder: a multicenter magnetic resonance imaging study. Archives of General Psychiatry, 69(2), 195–209. https://doi.org/10.1001/archgenpsychiatry.2011.1251

Ecker, C., Marquand, A., Mourão-Miranda, J., Johnston, P., Daly, E. M., Brammer, M. J., … & Murphy, D. G. (2010). Describing the brain in autism in five dimensions—magnetic resonance imaging-assisted diagnosis of autism spectrum disorder using a multiparameter classification approach. Journal of Neuroscience, 30(32), 10612–10623. https://doi.org/10.1523/JNEUROSCI.5413-09.2010

Eisenberg, I. W., Wallace, G. L., Kenworthy, L., Gotts, S. J., & Martin, A. (2015). Insistence on sameness relates to increased covariance of gray matter structure in autism spectrum disorder. Molecular Autism, 6, 54. https://doi.org/10.1186/s13229-015-0047-7

Fischl, B. (2012). FreeSurfer. NeuroImage, 62(2), 774–781. https://doi.org/10.1016/j.neuroimage.2012.01.021

Fishman, I., Keown, C. L., Lincoln, A. J., Pineda, J. A., & Müller, R.-A. (2014). Atypical Cross Talk Between Mentalizing and Mirror Neuron Networks in Autism Spectrum Disorder. In JAMA Psychiatry (Vol. 71, Issue 7, p. 751). https://doi.org/10.1001/jamapsychiatry.2014.83

Fitzgerald, J., Leemans, A., Kehoe, E., O’Hanlon, E., Gallagher, L., & McGrath, J. (2018). Abnormal fronto-parietal white matter organisation in the superior longitudinal fasciculus branches in autism spectrum disorders. In European Journal of Neuroscience (Vol. 47, Issue 6, pp. 652–661). https://doi.org/10.1111/ejn.13655

Frith, U., & Frith, C. (2001). The Biological Basis of Social Interaction. Current Directions in Psychological Science, 10(5), 151–155. https://doi.org/10.1111/1467-8721.00137

Girgis, R. R., Minshew, N. J., Melhem, N. M., Nutche, J. J., Keshavan, M. S., & Hardan, A. Y. (2007). Volumetric alterations of the orbitofrontal cortex in autism. Progress in Neuro-Psychopharmacology & Biological Psychiatry, 31(1), 41–45. https://doi.org/10.1016/j.pnpbp.2006.06.007

Groen, W. B., Buitelaar, J. K., van der Gaag, R. J., & Zwiers, M. P. (2011). Pervasive microstructural abnormalities in autism: a DTI study. Journal of Psychiatry & Neuroscience: JPN, 36(1), 32–40. https://doi.org/10.1503/jpn.090100

Hadjikhani, N., Joseph, R. M., Snyder, J., & Tager-Flusberg, H. (2006). Anatomical differences in the mirror neuron system and social cognition network in autism. Cerebral Cortex, 16(9), 1276–1282. https://doi.org/10.1093/cercor/bhj069

Hardan, A. Y., Girgis, R. R., Lacerda, A. L. T., Yorbik, O., Kilpatrick, M., Keshavan, M. S., & Minshew, N. J. (2006). Magnetic resonance imaging study of the orbitofrontal cortex in autism. Journal of Child Neurology, 21(10), 866–871. https://doi.org/10.1177/08830738060210100701

Hardan, A. Y., Minshew, N. J., Mallikarjuhn, M., & Keshavan, M. S. (2001). Brain Volume in Autism. In Journal Of Child Neurology (Vol. 16, Issue 06, p. 421). https://doi.org/10.2310/7010.2001.7113

Hollander, E., Anagnostou, E., Chaplin, W., Esposito, K., Haznedar, M. M., Licalzi, E., Wasserman, S., Soorya, L., & Buchsbaum, M. (2005). Striatal volume on magnetic resonance imaging and repetitive behaviors in autism. Biological Psychiatry, 58(3), 226–232. https://doi.org/10.1016/j.biopsych.2005.03.040

Hyde, K. L., Samson, F., Evans, A. C., & Mottron, L. (2010). Neuroanatomical differences in brain areas implicated in perceptual and other core features of autism revealed by cortical thickness analysis and voxel◻based morphometry. Human brain mapping, 31(4), 556–566. https://doi.org/10.1002/hbm.20887

Itahashi, T., Yamada, T., Nakamura, M., Watanabe, H., Yamagata, B., Jimbo, D., Shioda, S., Kuroda, M., Toriizuka, K., Kato, N., & Hashimoto, R. (2015). Linked alterations in gray and white matter morphology in adults with high-functioning autism spectrum disorder: A multimodal brain imaging study. In NeuroImage: Clinical (Vol. 7, pp. 155–169). https://doi.org/10.1016/j.nicl.2014.11.019

Jiao, Y., Chen, R., Ke, X., Chu, K., Lu, Z., & Herskovits, E. H. (2010). Predictive models of autism spectrum disorder based on brain regional cortical thickness. NeuroImage, 50(2), 589–599. https://doi.org/10.1016/j.neuroimage.2009.12.047

Keehn, B., Brenner, L., Palmer, E., Lincoln, A. J., & Müller, R.-A. (2008). Functional brain organization for visual search in ASD. Journal of the International Neuropsychological Society: JINS, 14(6), 990–1003. https://doi.org/10.1017/S1355617708081356

Knaus, T. A., Kamps, J., Foundas, A. L., & Tager-Flusberg, H. (2018). Atypical PT anatomy in children with autism spectrum disorder with expressive language deficits. Brain Imaging and Behavior, 12(5), 1419–1430. https://doi.org/10.1007/s11682-017-9795-7

Langen, M., Leemans, A., Johnston, P., Ecker, C., Daly, E., Murphy, C. M., dell’Acqua, F., Durston, S., & Murphy, D. G. M. (2012). Fronto-striatal circuitry and inhibitory control in autism: Findings from diffusion tensor imaging tractography. In Cortex (Vol. 48, Issue 2, pp. 183–193). https://doi.org/10.1016/j.cortex.2011.05.018

Langen, M., Schnack, H. G., Nederveen, H., Bos, D., Lahuis, B. E., de Jonge, M. V., van Engeland, H., & Durston, S. (2009). Changes in the developmental trajectories of striatum in autism. Biological Psychiatry, 66(4), 327–333. https://doi.org/10.1016/j.biopsych.2009.03.017

Lee, J. E., Bigler, E. D., Alexander, A. L., Lazar, M., DuBray, M. B., Chung, M. K., Johnson, M., Morgan, J., Miller, J. N., McMahon, W. M., Lu, J., Jeong, E.-K., & Lainhart, J. E. (2007). Diffusion tensor imaging of white matter in the superior temporal gyrus and temporal stem in autism. In Neuroscience Letters (Vol. 424, Issue 2, pp. 127–132). https://doi.org/10.1016/j.neulet.2007.07.042

Lewis, P. A., Rezaie, R., Brown, R., Roberts, N., & Dunbar, R. I. M. (2011). Ventromedial prefrontal volume predicts understanding of others and social network size. NeuroImage, 57(4), 1624–1629. https://doi.org/10.1016/j.neuroimage.2011.05.030

Libero, L. E., Schaer, M., Li, D. D., Amaral, D. G., & Nordahl, C. W. (2019). A Longitudinal Study of Local Gyrification Index in Young Boys With Autism Spectrum Disorder. Cerebral Cortex, 29(6), 2575–2587. https://doi.org/10.1093/cercor/bhy126

Lisiecka, D. M., Holt, R., Tait, R., Ford, M., Lai, M.-C., Chura, L. R., Baron-Cohen, S., Spencer, M. D., & Suckling, J. (2015). Developmental white matter microstructure in autism phenotype and corresponding endophenotype during adolescence. Translational Psychiatry, 5, e529. https://doi.org/10.1038/tp.2015.23

Mahajan, R., Dirlikov, B., Crocetti, D., & Mostofsky, S. H. (2016). Motor Circuit Anatomy in Children with Autism Spectrum Disorder With or Without Attention Deficit Hyperactivity Disorder. Autism Research: Official Journal of the International Society for Autism Research, 9(1), 67–81. https://doi.org/10.1002/aur.1497

Mueller, S., Keeser, D., Samson, A. C., Kirsch, V., Blautzik, J., Grothe, M., Erat, O., Hegenloh, M., Coates, U., Reiser, M. F., Hennig-Fast, K., & Meindl, T. (2013). Convergent Findings of Altered Functional and Structural Brain Connectivity in Individuals with High Functioning Autism: A Multimodal MRI Study. PloS One, 8(6), e67329. https://doi.org/10.1371/journal.pone.0067329

Mundy, P. (2003). Annotation: The neural basis of social impairments in autism: The role of the dorsal medial-frontal cortex and anterior cingulate system. Journal of Child Psychology and Psychiatry, and Allied Disciplines, 44(6), 793–809.

Mundy, P. (2018). A review of joint attention and social-cognitive brain systems in typical development and autism spectrum disorder. The European Journal of Neuroscience, 47(6), 497–514. https://doi.org/10.1111/ejn.13720

Nebel, M. B., Eloyan, A., Barber, A. D., & Mostofsky, S. H. (2014). Precentral gyrus functional connectivity signatures of autism. Frontiers in Systems Neuroscience, 8, 80. https://doi.org/10.3389/fnsys.2014.00080

Nickl◻Jockschat, T., Habel, U., Maria Michel, T., Manning, J., Laird, A. R., Fox, P. T., … & Eickhoff, S. B. (2012). Brain structure anomalies in autism spectrum disorder—A meta◻analysis of VBM studies using anatomic likelihood estimation. Human brain mapping, 33(6), 1470–1489. DOI: 10.1002/hbm.21299

Pappaianni, E., Siugzdaite, R., Vettori, S., Venuti, P., Job, R., & Grecucci, A. (2018). Three shades of grey: detecting brain abnormalities in children with autism using source◻, voxel◻and surface◻based morphometry. European Journal of Neuroscience, 47(6), 690–700. https://doi.org/10.1111/ejn.13704

Pelphrey, K. A., Shultz, S., Hudac, C. M., & Vander Wyk, B. C. (2011). Research review: Constraining heterogeneity: the social brain and its development in autism spectrum disorder. Journal of Child Psychology and Psychiatry, and Allied Disciplines, 52(6), 631–644. https://doi.org/10.1111/j.1469-7610.2010.02349.x

Peterson, E., Schmidt, G. L., Tregellas, J. R., Winterrowd, E., Kopelioff, L., Hepburn, S., Reite, M., & Rojas, D. C. (2006). A voxel-based morphometry study of gray matter in parents of children with autism. Neuroreport, 17(12), 1289–1292. https://doi.org/10.1097/01.wnr.0000233087.15710.87

Raven, J. (2000). The Raven’s progressive matrices: change and stability over culture and time. Cognitive Psychology, 41(1), 1–48. https://doi.org/10.1006/cogp.1999.0735

Rippon, G., Brock, J., Brown, C., & Boucher, J. (2007). Disordered connectivity in the autistic brain: challenges for the “new psychophysiology.” International Journal of Psychophysiology: Official Journal of the International Organization of Psychophysiology, 63(2), 164–172.

Robinson, E. B., Koenen, K. C., McCormick, M. C., Munir, K., Hallett, V., Happé, F., Plomin, R., & Ronald, A. (2011). Evidence that autistic traits show the same etiology in the general population and at the quantitative extremes (5%, 2.5%, and 1%). Archives of General Psychiatry, 68(11), 1113–1121. https://doi.org/10.1001/archgenpsychiatry.2011.119

Rojas, D. C., Peterson, E., Winterrowd, E., Reite, M. L., Rogers, S. J., & Tregellas, J. R. (2006). Regional gray matter volumetric changes in autism associated with social and repetitive behavior symptoms. BMC Psychiatry, 6, 56. https://doi.org/10.1186/1471-244X-6-56

Sabbagh, M. A. (2004). Understanding orbitofrontal contributions to theory-of-mind reasoning: implications for autism. Brain and Cognition, 55(1), 209–219. https://doi.org/10.1016/j.bandc.2003.04.002

Sato, W., Kochiyama, T., Uono, S., Yoshimura, S., Kubota, Y., Sawada, R., Sakihama, M., & Toichi, M. (2017). Reduced Gray Matter Volume in the Social Brain Network in Adults with Autism Spectrum Disorder. Frontiers in Human Neuroscience, 11, 395. https://doi.org/10.3389/fnhum.2017.00395

Sato, W., Kubota, Y., Kochiyama, T., Uono, S., Yoshimura, S., Sawada, R., Sakihama, M., & Toichi, M. (2014). Increased putamen volume in adults with autism spectrum disorder. Frontiers in Human Neuroscience, 8, 957. https://doi.org/10.3389/fnhum.2014.00957

Schuetze, M., Park, M. T. M., Cho, I. Y., MacMaster, F. P., Chakravarty, M. M., & Bray, S. L. (2016). Morphological Alterations in the Thalamus, Striatum, and Pallidum in Autism Spectrum Disorder. Neuropsychopharmacology: Official Publication of the American College of Neuropsychopharmacology, 41(11), 2627–2637. https://doi.org/10.1038/npp.2016.64

Schultz, R. T. (2005). Developmental deficits in social perception in autism: the role of the amygdala and fusiform face area. International Journal of Developmental Neuroscience: The Official Journal of the International Society for Developmental Neuroscience, 23(2-3), 125–141. https://doi.org/10.1016/j.ijdevneu.2004.12.012

Shukla, D. K., Keehn, B., & Müller, R. A. (2011). Tract◻specific analyses of diffusion tensor imaging show widespread white matter compromise in autism spectrum disorder. Journal of Child Psychology and Psychiatry, and Allied Disciplines.

Smith, S. M., Jenkinson, M., Johansen-Berg, H., Rueckert, D., Nichols, T. E., Mackay, C. E., Watkins, K. E., Ciccarelli, O., Zaheer Cader, M., Matthews, P. M., & Behrens, T. E. J. (2006). Tract-based spatial statistics: Voxelwise analysis of multi-subject diffusion data. In NeuroImage (Vol. 31, Issue 4, pp. 1487–1505). https://doi.org/10.1016/j.neuroimage.2006.02.024

Valk, S. L., Di Martino, A., Milham, M. P., & Bernhardt, B. C. (2015). Multicenter mapping of structural network alterations in autism. Human brain mapping, 36(6), 2364–2373. https://doi.org/10.1002/hbm.22776

Via, E., Radua, J., Cardoner, N., & Happé, F. (2011). Meta-analysis of gray matter abnormalities in autism spectrum disorder: should Asperger disorder be subsumed under a broader umbrella of autistic spectrum …. Archives of General Psychiatry. https://jamanetwork.com/journals/jamapsychiatry/article-abstract/211201

Wass, S. (2011). Distortions and disconnections: disrupted brain connectivity in autism. Brain and Cognition, 75(1), 18–28. https://doi.org/10.1016/j.bandc.2010.10.005

Whitehouse, A. J. O., Hickey, M., & Ronald, A. (2011). Are autistic traits in the general population stable across development? PloS One, 6(8), e23029. https://doi.org/10.1371/journal.pone.0023029

